# Rapid and affordable generation of compensation beads for nanoscale flow cytometry

**DOI:** 10.64898/2026.04.24.720194

**Authors:** Johann Mar Gudbergsson, Anders Etzerodt

**Affiliations:** Department of Biomedicine, Aarhus University, Aarhus, Denmark

**Author notes:** Correspondence, Johann Mar Gudbergsson, Ph.D.

**Keywords:** nano flow cytometry, nanoflow, conjugation, antibody, fluorochrome, polystyrene beads

## Abstract

With the introduction of dedicated nanoscale flow cytometers, the need for suitable compensation beads has emerged. Here, we present a rapid and cost-effective method to generate ∼100 nm antibody-binding compensation beads compatible with a wide range of antibody species for use in nanoscale flow cytometry. This approach may provide a practical interim solution until commercial alternatives become available.

## Materials and methods

### Materials

- 250 mM borate buffered saline pH 8.5 (5x solution), filter sterilized 0.2 um
- 1M Tris-HCl pH pH 8.5, filter sterilized 0.2 um
- Sterile ddH_2_O
- Syringe filters 0.2 um
- Syringe 3 mL
- Recombinant protein A/G (Cat. #ab52213, Abcam)
- 100 nm polystyrene beads, NHS-functionalized (Cat. #ABVI-ABPS-0010-NHS-5ml, Abvigen)
- Fluorochrome-conjugated antibodies (See table 1 for antibodies used in this study)

**Table 1.** Detailed overview of antibodies used.

| Fluorochrome | Species, subclass | Clone | Catalogue | Stock concentration mg/ml | Applied ab vol/ mass per sample |
| --- | --- | --- | --- | --- | --- |
| BV421 | Ar. Ha. Igg2 | 145-2C11 | #562600, BD Horizon | 0.2 | 0.0625 ug - 1 ug |
| APC | Rat igg1 | 9-4D2-1E4 | #347305, BioLegend | N/A | 0.5 ul |
| APC | Rat igg2a | NVG-2 | #143906, BioLegend | 0.2 | 0.5 ul |
| APC | Ms igg1 | X54-5/7.1 | #139305, BioLegend | 0.2 | 0.5 ul |
| APC | Ms igg2a | 104 | #109813, BioLegend | 0.2 | 0.5 ul |
| APC | Hu igg1 | REA563 | #130-108-924, Miltenyi | 0.15 | 0.5 ul |
| APC | Ar. Ha. igg1 | Eat-2 | #104909, BioLegend | 0.2 | 0.5 ul |
| FITC | Rat igg2a | TNKUPJ | #11-1631-82, eBioscience | 0.5 | 0.5 ul |
| AF488 | Rat igg1 | ALY7 | #53-0443-82, Invitrogen | 0.5 | 0.5 ul |
| PE | Rat igg2a | MZ3 | #124805, BioLegend | 0.2 | 0.5 ul, 1.25 ul |
| AF647 | Rat igg2a | MZ3 | #124809, BioLegend | 0.5 | 0.5 ul |

Do not use autoclaved tubes as they might add unnecessary background particles, as we have previously shown [1].

### Instrument

CytoFLEX Nano

### Preparing 250 mM borate buffered saline

1. To make 50 mL, weigh the following into an appropriate container:
  - 0.62g Boric acid (H_3_BO_3_)
  - 0.44g Sodium chloride (NaCl)
  - 0.5g Sodium tetraborate decahydrate (Na_2_B_4_O_7_ · 10H_2_O)
2. Add 40 mL of sterile ddH_2_O, place on a magnetic stirrer until fully dissolved.
3. Adjust pH to 8.5 using 5N NaOH.
4. Bring volume to 50 mL with sterile ddH_2_O.
5. Run through a pre-wetted 0.2 um filter, i.e. using a 3 mL syringe and a 0.2 um syringe filter.

### Coupling of recombinant Protein A/G to 100nm NHS-functionalized polystyrene beads

1. Protein A/G was dissolved in sterile ddH_2_O at a concentration of 5 mg/mL
2. 50 ug of Protein A/G was conjugated to 0.5 mg of NHS-beads in a 100 µL reaction
  - 50 µL water,
  - 20 µL beads
  - 20 µL borate buffered saline
  - 10 µL Protein A/G
3. Mix well by vortexing or handpipetting
4. Incubate at RT for 2 hours.
5. Quench reaction by adding 5 µL of 1M sterile filtered Tris-HCl.
6. Leave for 15-30 min. at RT before use.
7. Store at 4°C until use. DO NOT FREEZE.

### Preparation of antibody-bound compensation beads for nano flow cytometry

1. Dilute appropriate amount of compensation beads 1:125 in sterile ddH_2_O
  a. *The bead dilutions can be further optimized for more cost-effective use*.
2. Dilute 0.5 µL antibody in 25 µL sterile ddH_2_O
  a. *Antibody vials with low concentration and dim fluorochromes might need increased amount to saturate beads*.
3. Mix 25 µL of diluted beads with 25 µL of diluted antibody.
4. Incubate for 5-10 min. at RT protected from light
5. Dilute 1:80 in sterile ddH_2_O and load into the CytoFLEX Nano or other nanoscale flow cytometer.
6. Set vSSC1-H threshold (or comparative SSC threshold if using a different cytometer) to >500.
7. Keep fluorescence detector gains at QC levels unless fluorochrome-specific adjustments are needed.
8. Run at appropriate speed to between 400 – 1000 events per second.
  a. *In our case, 1 µL/min was sufficient for 400 events/s*
9. Acquire for 1 min.
10. Run all appropriate samples and controls and perform compensation.

## Results & discussion

Compensation beads for cell-oriented flow cytometry are often several microns in diameter which renders them too large for dedicated nanoscale flow cytometers. Here, we set out to generate nano compensation (nComp) beads for which we developed a simple protocol that can be implemented in any standard laboratory. Fortunately, we identified commercially available 100 nm NHS-functionalized polystyrene beads, making the protocol readily accessible and easy to adopt.

First, we demonstrated that the conjugation protocol did not result in measurable changes in the physical properties of the nano compensation (nComp) beads and that the event rate remained stable (Figure 1A). Using a BV421-conjugated antibody, we performed titrations at both QC fluorescence gain and maximum gain settings. All peaks were clearly separated from the blank, and a dose-dependent increase in signal was observed under both gain conditions (Figure 1B–C). To directly compare the gain settings, we calculated the stain index 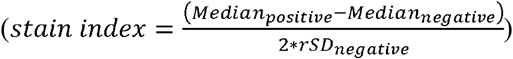 and found that both performed similarly (Figure 1C). Consequently, all subsequent experiments were carried out using the QC gain settings. The nComp beads bound multiple antibody species and subclasses, with human IgG1 showing the lowest binding and rat IgG2a the highest (Figure 1D). We also evaluated multiple fluorochromes across the four lasers and detected clear signals from nearly all of them (Figure 1E). Although PE is one of the brightest fluorochromes, the antibody conjugated with PE showed only a weak separation from the blank compared to other fluorochromes. To increase staining intensity, we increased the concentration of PE-conjugated antibody which resulted in an improved separation (Figure 1E). As nComp beads are only ∼100nm and PE is one of the largest fluorochromes used in flow cytometry, a possible explanation for the reduced sensitivity of PE-conjugated antibodies could be steric hindrance. Interestingly, FITC-conjugated antibodes also showed a strikingly low separation from the blank. However, previously published results suggest that using pure water as a diluent might affect the quantum yield and consequently reduces brightness [2], [3]. Thus, testing different dilution buffers could be a solution in case you encounter a fluorochrome that is sensitive to the buffer composition.

**Figure 1.**
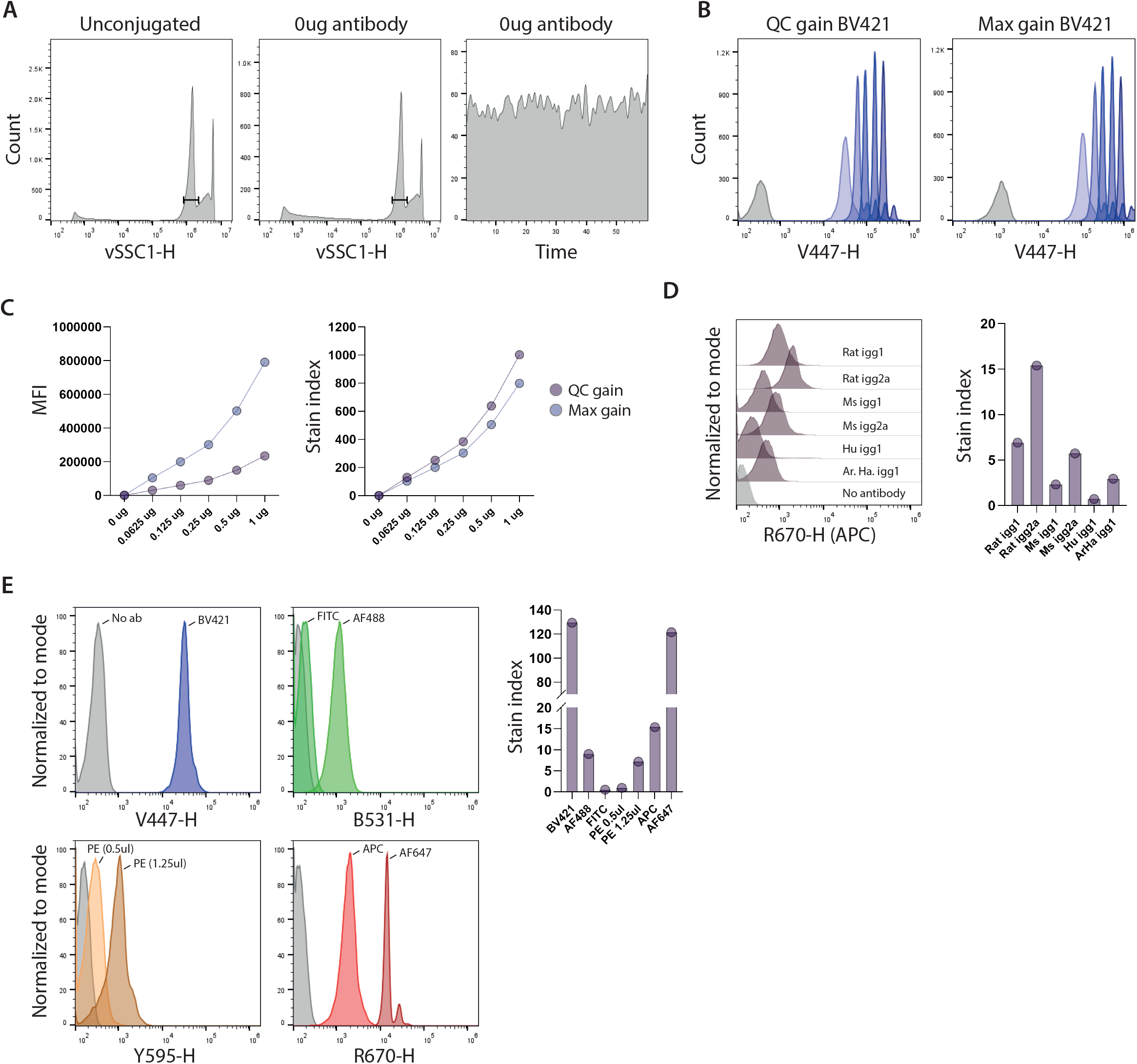

## Concluding remarks

In summary, we have successfully generated nComp beads suitable as compensation controls for experiments performed on dedicated nanoscale flow cytometers, such as the CytoFLEX Nano. As no suitable commercial compensation beads are currently available, we anticipate that this approach will enable researchers to produce their own reagents until such alternatives become accessible. Optimal dilutions and antibody concentrations should be determined by individual laboratories, and the shelf life of the conjugated beads remains unknown; therefore, regular quality control may be required.

## Acknowledgement

The authors wish to acknowledge the FACS Core Facility at Aarhus University. The CytoFLEX Nano was a generous gift from the Carlsberg Foundation (Grant CF23-1020).

